# Curiosity shapes brain-like architectures and functions

**DOI:** 10.64898/2026.07.02.735826

**Authors:** Francesco Poli, Kayson Fakhar, Alexa Mousley, Cheslie C. Klein, Charlotte Grosse Wiesmann, Duncan E. Astle

## Abstract

How does complex cognition emerge from simpler underlying processes? We show that two components are sufficient: infant-like curiosity and brain-like biophysical constraints *jointly* drive the emergence of complex neuronal architectures and cognitive abilities. We first tested curiosity-driven exploration in 275 8-to 15-month-old infants. We then implemented these mechanisms in artificial recurrent neural networks, letting them actively sample 20 cognitive tasks during training, while forcing the networks to operate under brain-like biophysical constraints. These two components were sufficient for a range of biological and cognitive phenomena to emerge: The resulting networks captured human synaptic development, the adult brain’s architecture, and displayed compositional generalisation, a hallmark of human cognition. Curiosity, long viewed as a downstream consequence of complex brains, is also a driver of their complexity.

---

The human mind builds an unbounded range of thoughts, plans, and inferences, recombining simpler elements into endlessly new configurations. More than a century of inquiry has given us extraordinarily detailed descriptions of the behavioural signatures^1,2^, computational mechanisms^3–5^, and neural correlates^6–8^ of this complex cognition. And yet the question of how that complexity actually *arises* has stayed out of reach: by what mechanism does a system with simple parts become a mind capable of such complex cognition? Some developmental theories have argued for decades that cognition is not prespecified but self-organises out of biological constraints interacting over developmental time^9–11^. However, the claim has been hard to test directly: doing so would require ablating and recombining biological constraints during development and observing the cognitive outcome, an intervention no developing brain allows.

Here, we propose a different approach: equipping artificial recurrent neural networks (RNNs) with a small set of fundamental cognitive and biological components, and asking whether complex cognition can emerge as a result. Within a neuroconnectionist research programme^12^, RNNs have been used widely to understand cognitive abilities and their underlying neural processes^8,13–18^. We use them here to probe the minimal requirements for the emergence of complex cognition. To this end, we equipped populations of RNNs with two key human-like components: curiosity-driven exploration and brain-like biophysical constraints. The networks shared a minimal common architecture (an input layer encoding task identity, a fixation signal, and sensory stimuli; a single recurrent hidden layer of 100 units; and an output layer producing the motor response; see Fig. 1A) and were trained on a battery of tasks typically used to test higher-order cognition in humans (Fig. 1C–D).

**Fig. 1.**
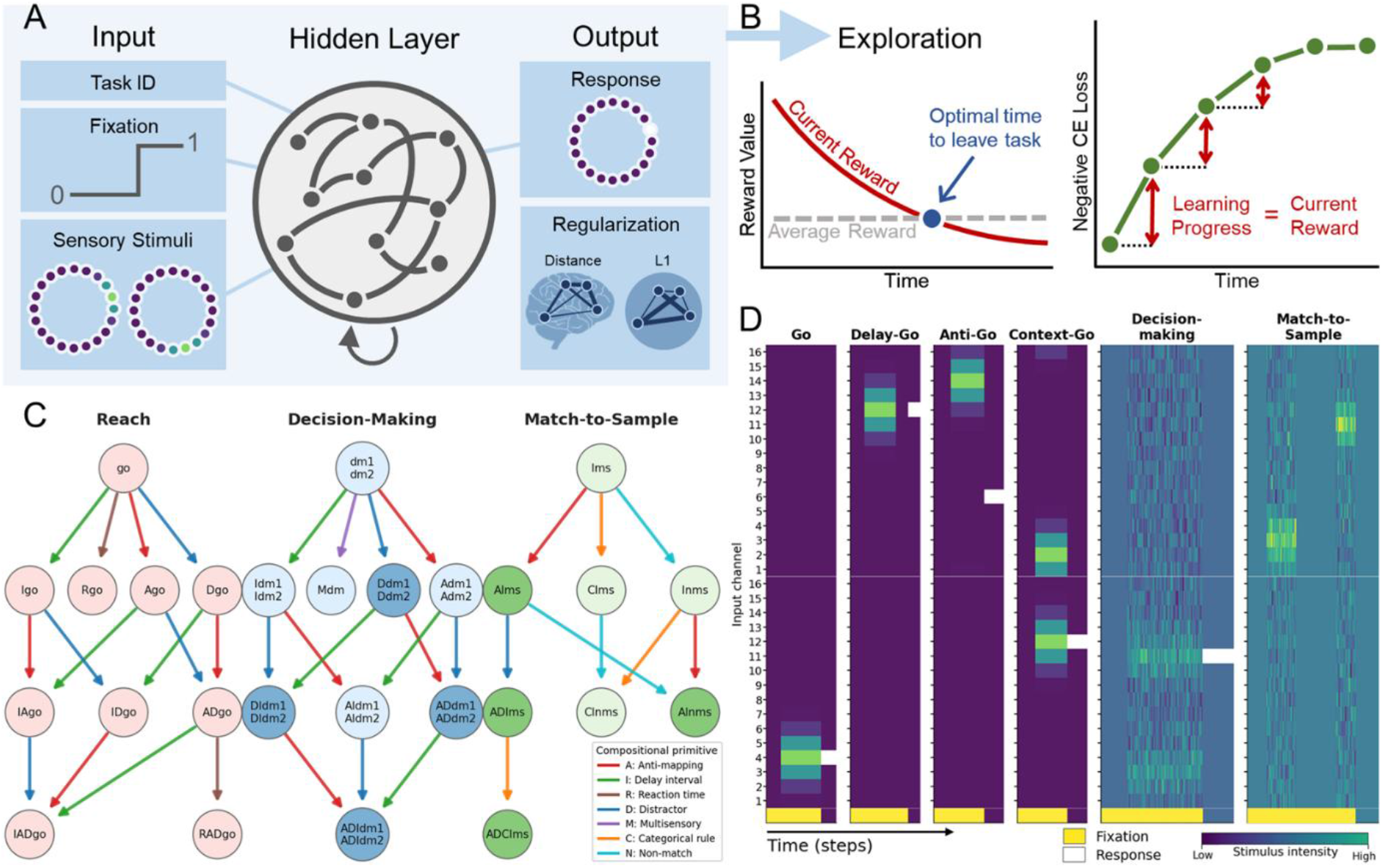
Learning multiple tasks through curiosity-driven exploration. **A**. RNN with task-identity, fixation, and ring-coded sensory inputs; a single recurrent hidden layer of 100 units; and a motor-response output. Recurrent weights are regularised across the entire duration of the training by L1 or by a distance penalty over Schaefer-100 coordinates. **B**. Task selection during training follows the marginal value theorem: the network leaves a task when its current learning progress (reduction in cross-entropy loss) drops below the running global average. **C**. Compositional task set. Three families of tasks were generated by combining a base task (Reach, Decision-Making, Match-to-Sample) with primitive modifiers (Anti, Delay, Reaction-time, Distractor, Multisensory, Categorical, Non-match). RNNs were trained on 20 tasks; 12 more tasks were held out for test. **D**. Trial time course for six example tasks; yellow bars mark fixation, white intervals the correct response.

## Infant-like curiosity and biophysical constraints in recurrent neural networks

The first human-like component we implemented in RNNs reflects a long-standing principle that proactive sampling of the world is formative for cognition^10,19^. Curiosity-driven exploration is simply the more recent, computational operationalisation of this principle, providing finer-grained insight into the mechanisms by which humans actively sample the world across the lifespan^20,21^. The second component draws on a more recent tradition in network neuroscience showing that the biophysical constraints faced by the brain, such as the cost of long-range connections and the geometric arrangement of regions, substantially determine the computations a brain can support^22–24^. We address each in turn.

Curiosity-driven exploration has been instrumental in robotics^25–27^ and, more recently, in artificial intelligence^28–31^, where it lets agents learn in environments without explicit external reward. However, the mechanisms endowed in these models (e.g., exhaustively sampling every option before committing to one) are biologically implausible. Infants, by contrast, must decide when to explore from a limited set of past experiences alone. Implementing infant-like curiosity in RNNs is challenging, because the very mechanisms underlying it remain debated^32–34^.

To address this, we first collected eye-tracking data from 275 infants on a visual paradigm already established to capture curiosity-driven sampling^33,35,36^. Stimuli appeared at four different locations on the screen following probabilistic, learnable sequences (Fig. 2D). Stimuli differed in how informative they were to learn the underlying probabilistic structure of the sequences. We quantified the information content of each stimulus precisely using Kullback-Leibler divergence^37^. This measure captures how much new information each stimulus brings, compared to the information contained in all the preceding stimuli in the sequence. While the stimuli were presented, we tracked the infants’ trial-by-trial decisions to either stay engaged with the stimuli (i.e., keep looking) or disengage (i.e., look away).

**Figure 2.**
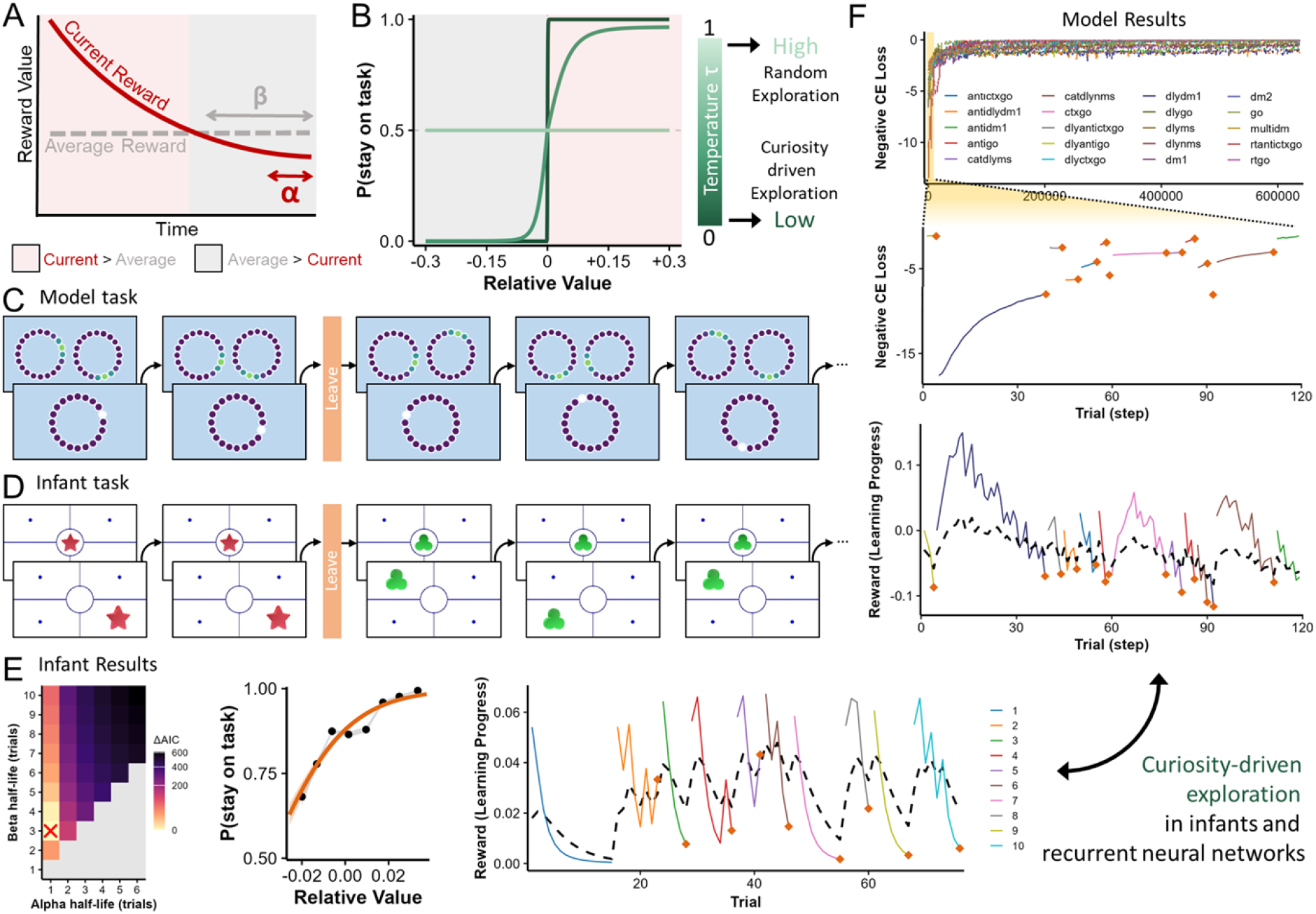
Infant-like curiosity in RNNs. **A**. The reward of the current task is tracked by a fast moving average with rate **α**; a slower moving average with rate **β** tracks the average reward across longer timescales. **B**. The agent leaves a task when its current reward falls below the running average. This decision is stochastic, regulated by a temperature parameter **τ**. Low τ produces deterministic, reward-driven task switching; high τ makes task switching essentially random. **C-D**. Model and infant tasks share the same foraging structure: the agent can leave at any moment for an alternative. **E**. Infant results. *Left*: ΔAIC across an (α, β) half-life grid; the red × marks the best fit. *Middle*: fitted MVT sigmoid across all infants. *Right*: example trial sequence showing learning progress over time, with leaving events marked (orange diamonds; dashed line = running average). **F**. Example model performance. *Top:* Full training trajectory across all tasks. *Middle*: a zoomed-in segment. Bottom: the per-trial reward signal with leaving events marked (bottom).

We treated infants’ looking behavior as a series of foraging decisions. Differently from foraging tasks, here the value foraged is not extrinsic (e.g., food) but intrinsic: the opportunity to learn. This reformulation afforded the application of a well-established framework from foraging ecology, the marginal value theorem^38,39^, to capture curiosity-driven exploration (Fig. 2A). The marginal value theorem prescribes that an agent should abandon its current source of reward when it falls below the average reward. Across the new data and two pre-existing datasets, the marginal value theorem captured infants’ behaviour very well (*β* = −0.80, *SE* = 0.03, *z* = −25.1, *p* < 0.001, Fig. 2E) and more accurately than any prior formalisation^33,34^ (ΔAIC ≥ 414).

Once the mechanisms underlying curiosity-driven exploration were precisely formalised and identified in infants, we implemented them directly in RNNs. We generated populations of networks that differed in the extent to which their training relied on curiosity, regulated by a temperature parameter τ (Fig. 2B). Just like in infants, who could decide when to look away but not which sequence came next, curiosity determined when a network left its current task, not which task it moved to. At low τ, the decision to stay or leave followed the marginal value theorem deterministically, and was thus curiosity-driven; at high τ, it was effectively random; a degenerate regime that, incidentally, approximates the classical RNN training paradigm, in which tasks are interleaved as uniformly as possible. We also varied several other parameters governing the calibration of the marginal value theorem; none affected the main results (see Methods). The resulting RNNs reproduced infant-like exploration dynamics during training (Fig. 2F).

The second human-like component we implemented in RNNs is brain-like biophysical constraints. Implementing such constraints in RNNs is not straightforward, since RNNs are typically trained to maximise task performance and nothing else. A recent class of architectures (spatially embedded RNNs) was developed for precisely this purpose^13^. The key manipulation is to embed the network’s units in a physical 3D space and penalise each connection in proportion to the distance between the units it links, mimicking the wiring cost that makes long-range axons rare in real brains^40^. Here, we push this approach further by embedding the network not in an arbitrary 3D space but in the same one occupied by the brain: we used the coordinates of the Schaefer-100 parcellation and mapped them onto the 100 units of the RNN’s hidden layer, situating the network in human anatomical space. The population of RNNs that we generated varied systematically in regularisation strength used during training, with higher values producing stronger spatial penalisation. We also ran control RNNs that used classical L1 regularisation in place of the distance-based penalty.

Because the RNNs were embedded in the same anatomical space as the human brain, their connectomes could be directly compared in terms of network topology (Fig 3A). We focused on three signature properties widely used to characterise biological brains^41^: modularity, global efficiency, and rich-club organisation (Fig. 3B). For the human reference, we used the Human Connectome Project Young Adult (HCP-YA) dataset, comprising 1065 healthy individuals aged 22-35 years^42^; we parcellated each connectome with the Schaefer-100 atlas, extracted streamline counts, and pruned all networks to 13.5% density so that artificial and biological connectomes were comparable in edge count (the full preprocessing pipeline can be found in the Supplementary Information).

**Figure 3.**
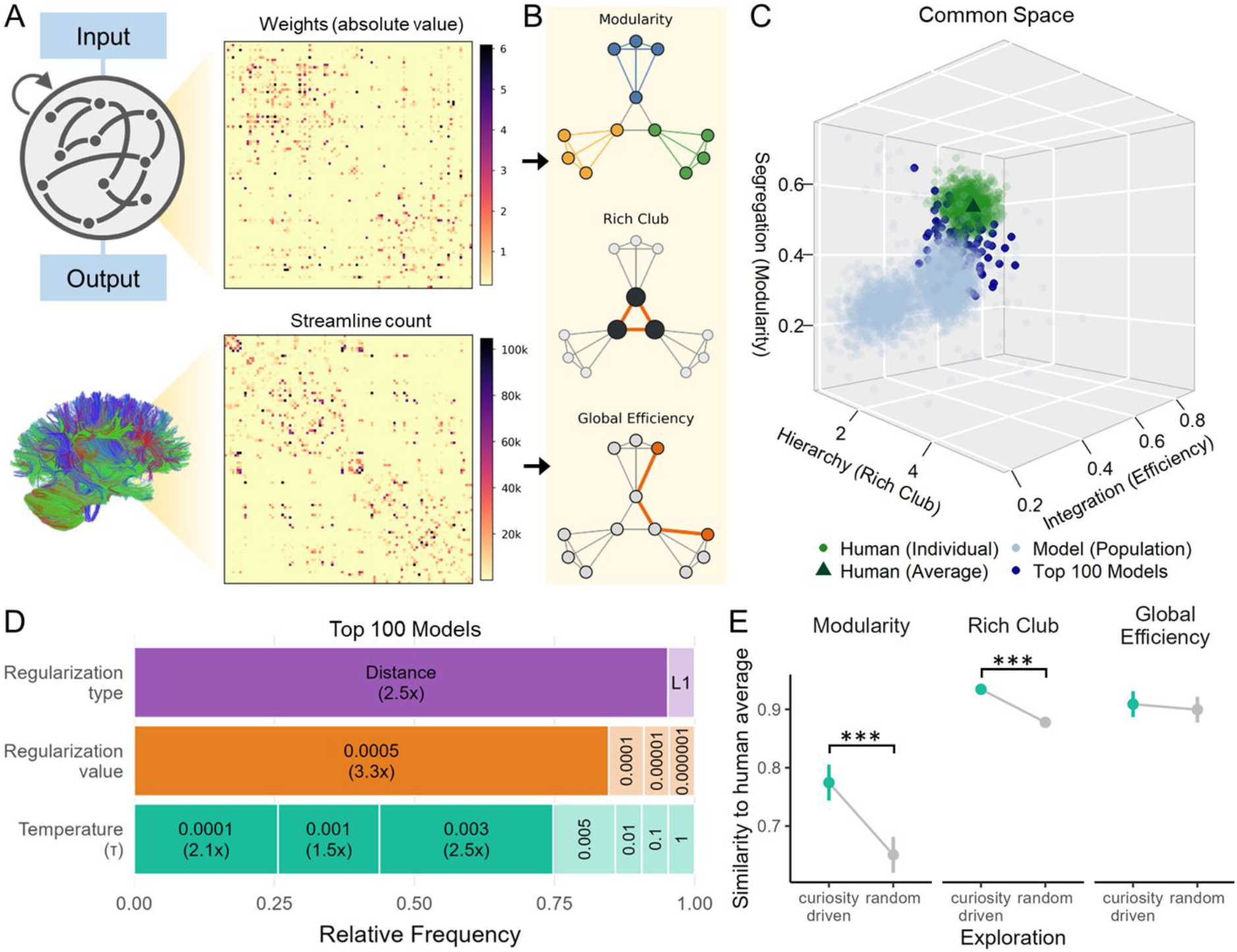
Network structure of RNNs and the human brain. **A**. Connectivity matrices used to compare the two systems. *Top*: absolute recurrent weights of an example RNN hidden layer. *Bottom*: streamline counts of an example human connectome reconstructed from white-matter tractography. **B**. Three signature topological properties computed identically on artificial and biological networks: *modularity* (segregation into densely connected sub-communities), *rich-club organisation* (presence of a densely interconnected hub backbone), and *global efficiency* (typical ease of communication between any two nodes). **C**. Common 3D space defined by these three metrics. Human brains (green) occupy a region characterised by high modularity, high efficiency, and a strong rich club; the 100 RNNs closest to the human average are highlighted in dark blue. **D**. Among the top 100 most human-like RNNs, distance regularisation, strong regularisation, and low τ are all over-represented. **E**. Comparing curiosity-driven and random-exploration RNNs shows that curiosity significantly increases modularity and rich-club organization, making the models’ hidden layer more similar to the human brain architecture.

### Emergence of complex architectures and functions in human-like RNNs

With both kinds of network placed in the same common space (Fig 3C), we could address the first central question: which of the components imposed on our RNNs were responsible for producing human-like brain networks? We first tested whether regularization type (distance-based vs L1), regularization value and temperature τ affected the RNNs’ similarity to human connectomes (quantified as a score s between 0 and 1, derived from the distance between each network’s topological fingerprint and the human average; see Methods).

As anticipated, distance-based regularisation and stronger regularisation value pulled networks toward biological topology (Δ*s* = 0.39, *t*_2368_ = 54.7, *p* < 0.0001), but, crucially, so did lower τ, particularly under strong distance-based regularisation (Δ*s* = 0.05, *t*_2368_ = 7.7, *p* < 0.0001). We confirmed this by identifying the top 100 RNNs whose connectomes were closest to the human reference and showing that strong distance-based regularization (*χ*^2^ = 145.5, *p* < 0.0001) and low temperature (*χ*^2^ = 76.6, *p* < 0.0001) were over-represented in this subset (Fig. 3D), indicating that the way a network samples its training experience is itself a determinant of how brain-like it becomes. Specifically, curiosity pushed RNNs towards more human-like levels of modularity (Δ*s* = 0.12, *t*_68_ = 5.7, *p* < 0.0001) and rich-club architecture (Δ*s* = 0.06, *t*_2368_ = 8.6, *p* < 0.0001) (Fig. 3E).

Having characterised network complexity at the end of training, we next examined how networks evolved over the course of training. For RNNs, we focused on the density of network weights, while for humans we focused on synaptic density. Both quantities measure how densely a system is wired at a given moment, and the comparison is possible because RNN weights are an abstraction that does not map onto any single biological element: just as they were compared with connections between regions in the topological analysis, their overall density can here be compared with the density of synapses. In humans, synaptic density follows a well-characterised trajectory: a steep rise in synapse number during the first years of life (synaptogenesis), followed by a slower pruning phase that begins in childhood and persists into adulthood^43^. However, no integrated dataset providing comparable measurements across multiple cortical regions was available. We therefore compiled one by harmonising the classical Huttenlocher studies^43,44^, yielding a comprehensive resource of synaptic density (synapses per 100 μm^3^) across the visual, auditory, and prefrontal cortices, spanning the fetal period through late adulthood (Fig. 4A). A generalised additive model fit to this dataset captured both synaptogenesis and pruning robustly (*edf* = 3.64, *p* < 0.0001, 85.8% of deviance explained).

**Figure 4.**
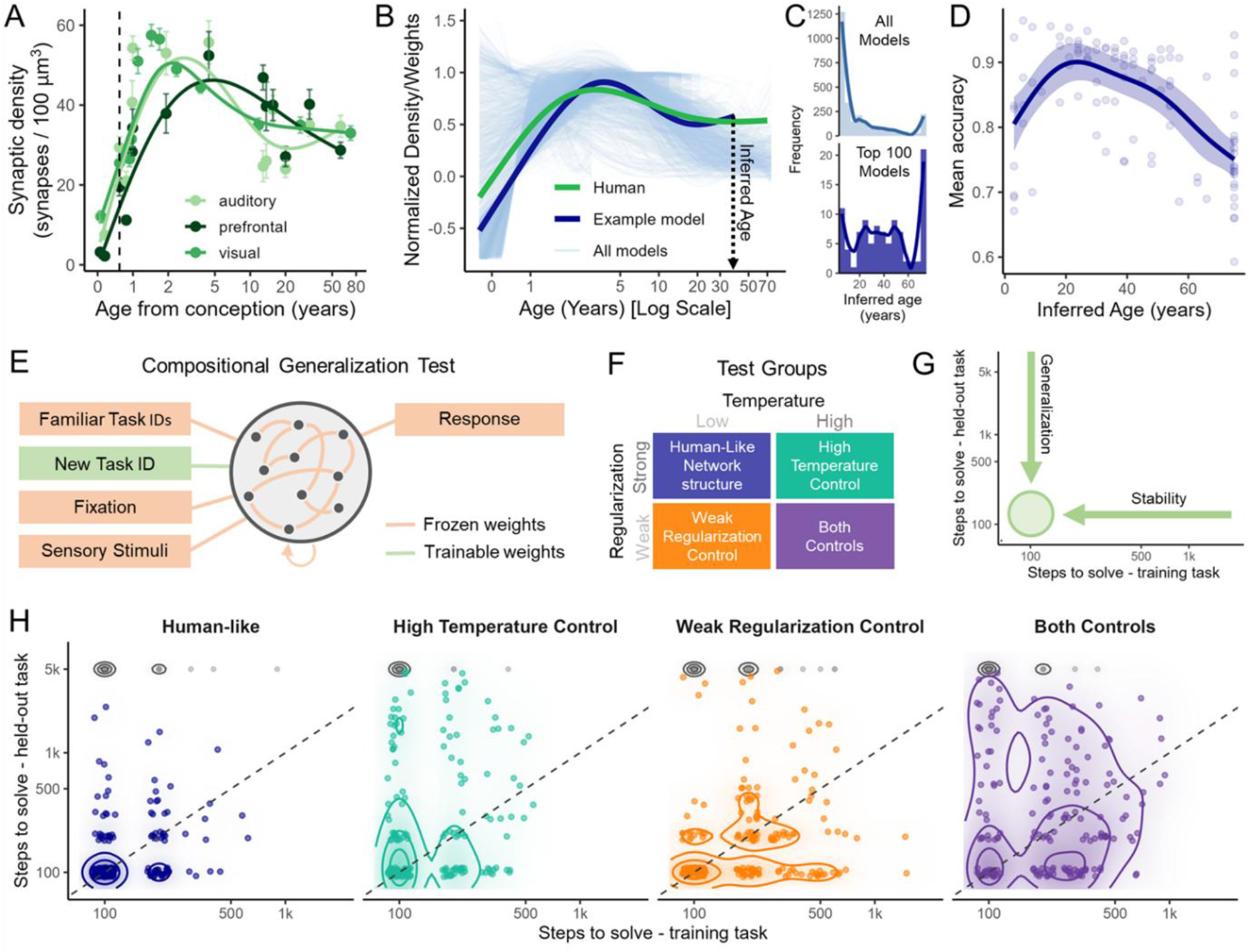
The emergence of complex cognition in human-like RNNs. **A**. Human synaptic density across the lifespan (mean ± SE; dashed line = birth; solid curves = Generalized Additive Models fit). **B**. Density trajectories for all RNNs and human data, with one example of how age was inferred for each model (dashed line). **C**. Distribution of inferred ages across all RNNs (top) and within the top 100 networks closest to the human reference in network topology (bottom). **D**. Mean task accuracy at the end of training as a function of inferred age. **E**. Compositional generalisation test. Every weight of the trained network is frozen (orange), and a new task identifier is introduced with fresh trainable connections into the hidden layer (green). **F**. Four matched cohorts crossed by temperature (Low/High) and regularisation value (Strong/Weak) were tested. **G**. Example of optimal performance in the compositional test: time to solve a training task (x) and time to solve its held-out compositional counterpart (y). Rapid resolution on the training axis reflects *stability* of internal dynamics; rapid resolution on the held-out axis reflects *compositional generalisation*. **H**. Joint distributions of training-task and held-out-task solution times for the four cohorts. Human-like networks concentrate in the lower-left corner, solving both training and held-out tasks quickly.

For RNNs, we tracked the total density of network weights across training steps. Weights change substantially during training because gradient descent strengthens connections that reduce task loss, while the spatial and L1 regularizers continually suppress those that contribute little. This produces highly variable trajectories of network density that can be compared with human synaptic density.

However, a direct comparison between RNNs’ and human synaptic density is challenging because their timescales (training steps vs biological age) are different. To bridge this gap, we treated training steps as the network’s developmental time. Then, we projected each RNN’s timescale onto the human lifespan by identifying the period of human development that most closely matched its density trajectory (Figure 4B). The match was evaluated on three properties of the model and human curves: their overall shape (correlation between the two curves), the timing of the peak relative to the lifespan, and how much density was pruned after the peak. The period that produced the closest match defined that RNN’s *fitted age*. This procedure gave every model a single fitted age, with substantial variability across the population.

Two findings followed. First, the 100 networks most similar to humans in topology also exhibited density trajectories closer to the human developmental arc than the rest of the networks (median fitted age 32 vs 9 years; *χ*^2^ = 88.7, *p* < 0.0001). Second, fitted age was a strong predictor of accuracy at the end of training (*edf* = 2.63, *F* = 15.9, *p* < 0.001, 36% of deviance explained): performance rose with fitted age up to roughly 20 years and declined thereafter, globally mirroring the rise and gradual decline of cognitive performance across the human lifespan^45^. Together, these results extend the previous finding: brain-like topology, brain-like developmental dynamics, and cognitive performance are not independent achievements but strongly co-vary, which is exactly the coupling expected if all three are downstream consequences of the same constraints.

Yet brain-like structure and developmental trajectory are not, on their own, what makes a mind: the decisive test is whether the same constraints also yield the hallmark of complex cognition. To test this, we compared four groups of networks on a compositional test (Fig. 4F). The human-like group comprised networks trained with curiosity and strong regularisation that were close to human topology and had learned the training tasks well. Three matched control groups were identical to them except that they explored at random (high τ), were only minimally regularised, or both (see Methods). These trained models were tested on tasks they had never encountered during training. These held-out tasks were constructed by recombining familiar task elements in new ways (e.g., composing the *distractor* and *decision-making* primitives, each of which the network had seen separately but never together).

Solving such tasks demands compositional generalization: the ability to recombine what has been learned into configurations never explicitly trained^46,47^. This complex cognitive ability is a hallmark of human cognition^48^ and one that artificial systems struggle to display^49^. To test this, we froze every weight of the trained networks and added a single new task identity with fresh trainable connections into the hidden layer. These new connections were the only weights allowed to change, and the networks could thus not learn any new internal dynamics. Under these conditions, if a held-out task is nevertheless solved quickly, it must be because the network dynamics already possessed the relevant primitives that support flexible recombination. As a control, we also tested every training task with the same approach: Instead of using the same task identity we used during training, we used a new one. This way, the network was blind to the fact that it had ever been trained on those tasks. Rapid re-solution of these familiar tasks indicates that the internal dynamics for known tasks are stable and reusable.

Results showed that human-like networks displayed both greater stability and stronger compositional generalisation, as evidenced by their faster resolution of training and held-out tasks alike (*χ*^2^ = 1071.0, *p* < 0.0001). This demonstrates that the same biological constraints that yield brain-like topology and brain-like developmental dynamics (i.e., curiosity and biophysical constraints) also produce a distinctive signature of complex cognition: the capacity to recombine learned primitives into configurations never explicitly trained. Hence, these findings indicate that curiosity-driven exploration and brain-like spatial constraints are sufficient for a key hallmark of complex cognition to emerge.

## Discussion

The claim that cognition self-organises from a small set of interacting constraints has remained largely conceptual. Testing it would require systematic ablation and recombination of cognitive and biological components, which cannot be performed in real brains. Here, we used artificial recurrent neural networks as a tractable experimental system to test this hypothesis. We equipped networks with two key components, curiosity-driven exploration and brain-like spatial organisation; these were sufficient for a key signature of human complex cognition to emerge alongside brain-like topology and developmental dynamics.

The contribution of curiosity here is important: how a network sampled its training experience proved to be a *determinant* of its final architecture, not merely a downstream consequence of it. At the mechanistic level, curiosity may pace a curriculum matched to the agent’s current competence, allowing the network to specialise gradually. Under biophysical pressure to segregate, this staged specialisation may resolve into meaningful, reusable and interconnected modules, a form of interactive specialisation also seen in infant development^50^. These results are also in line with two recent proposals about brain evolution: that the brain is a foraging device^39,51,52^, and that increased modularity is a defining feature of the human connectome’s evolution^53^. Our findings link the two, showing that foraging-like exploration *actively* shapes the modular organisation characteristic of the human brain.

Our human-like networks also showed better compositional generalisation: when frozen and probed on tasks that recombined familiar primitives in novel ways, they solved these held-out tasks rapidly, indicating that their internal dynamics already implemented the relevant primitives in a form that supported flexible recombination. The same networks even reproduced plausible trajectories of human synaptic and cognitive development, mirroring the rise and decline of cognitive performance across the lifespan. Together, these findings show that brain-like topology, brain-like development, and the hallmark of complex cognition are not independent achievements but coupled outcomes of *the same underlying constraints*. This accords with neuroconstructivist and dynamical-systems accounts of development, which hold that adult cognition arises from the interplay of brain maturation, experience, and the dynamical reorganisation of cognitive functions over time^9–11^. Our results give that long-standing theoretical view direct support at the level of whole-brain organization.

It is important to note that our model is a general system that does not attempt to capture the precise biology or psychology of the human brain, as it consists of a single hidden layer and was trained only on 20 tasks, whereas humans possess much more complex brains and learn many more skills across multiple domains. Nonetheless, using simple architectures on a small set of tasks is an approach has been proven fruitful in the past^16,17,54–57^. By manipulating only the components most relevant to our question (a curiosity algorithm, a flexible training curriculum and biophysical constraints), we can be more confident that our results are likely to reflect general principles of how neural systems develop rather than features of a particular architecture.

At the same time, these results open several avenues that were previously out of reach. Because any factor hypothesised to shape brain and cognitive development can be instantiated as an explicit constraint, the role of genetic, biological, cognitive, and environmental factors can be directly tested within our framework. Embodiment is the most immediate example: the human brain develops inside a body that dictates what can be sampled and which computations are worth performing^58^, and our approach can now test whether adding embodied constraints sharpens the emergence of complex brains and cognitive functions^59^. Beyond this, the models themselves generate testable predictions: they can be used to predict when and how specific cognitive capacities should emerge in human development, and these predictions can in turn be tested empirically. We see this as the beginning of a closer feedback loop between developmental theory, computational modelling, and behavioural neuroscience, which may reshape how we study the origins of complex cognition.

Finally, these findings have immediate relevance for research on artificial intelligence. The prevailing route to more capable systems is to make them ever larger, with rising computational, energetic, and economic costs^60^. Our results point the other way: human-like cognition arose not from scale^61^ but from the gradual unfolding of complexity under simple constraints, suggesting that more human-like, efficient, and interpretable systems may be reached by developing complexity rather than by enlarging models alone.

## Methods

### Infant Data

#### Participants

We pooled behavioral data from existing, previously published datasets^33,36,62^ and from a new data collection. For the existing data, a total of *N* = 173 infants were initially recruited at the Donders Institute for Cognition, Brain and Behavior (the Netherlands). After excluding 28 participants due to fussiness or technical error, a final *N* = 145 were retained (mean age = 8.19 months, SD = 0.62 months, Female = 47%). For the new data, infants were recruited at the Max Planck Institute for Human Cognitive and Brain Sciences (Germany). *N* = 131 infants were tested, of whom *N* = 130 were retained after the exclusion of one participant due to a technical error (mean age = 12.64 months, SD = 1.88 months, Female = 52%). All data was collected in accordance with local ethical guidelines after approval from the local ethical committee.

#### Apparatus and procedure

Infants were tested in a silent room with fixed light levels. Infants either sat in a baby seat held by the caregiver on their lap or directly on the caregiver’s lap, with their eyes approximately 60 cm from the screen. Looking behavior was monitored simultaneously with an eye-tracker (Tobii X300 or Tobii X120), and a video camera. Parents were instructed not to interact with their child unless the child explicitly sought their attention, and, in that case, not to try to redirect the child’s attention to the screen. Look-away events were coded from the video data using BORIS software^63^.

#### Visual learning task

Infants were presented with a visual learning task^33^ consisting of 16 sequences of cue-target trials. On each trial, a cue (i.e., a simple shape) appeared in the middle of the screen and was followed by a target (the same shape), at one of four quadrant locations around the cue. The shape was constant within a sequence and changed between sequences. Within each sequence, the target was more likely to appear at one specific location than at the other three; across the 16 sequences, the high-likelihood location received the target with probability 1.0 in 4 sequences, 0.8 in 6 sequences, and 0.6 in 6 sequences. The order of the 16 sequences was fixed across participants. A sequence contained at most 15 cue-target trials and terminated either when the infant had completed all 15 trials or when the infant looked away from the screen for at least one continuous second; in the latter case, the next sequence began as soon as the infant returned their gaze to the screen. The experiment ended when the infant had watched all 16 sequences or became fussy.

#### Look-away measure

For each target presentation we recorded a binary look-away event indicating whether the infant continued to attend to the screen or instead disengaged from it within the following 4 s (i.e., before the start of the next trial). We treat this event as an index of the infant’s current interest in the ongoing sequence^64^: because the testing setting contained no external distractors competing for the infant’s attention, a look-away reflects a self-initiated, active decision to stop engaging with the task^34^.

#### Computational Modeling

We modeled infants’ decisions to look away as foraging decisions under the Marginal Value Theorem (MVT)^38^. While MVT was originally formulated in behavioral ecology to capture the principle that an animal abandons a depleting food patch when its local return falls below the average return available in the environment, its logic extends naturally to any setting in which an agent must allocate time across discrete sources of value. Here, we treat each sequence as a patch in which the infant harvests learning progress, and treat look-aways as decisions to leave the current patch in favor of another one. Curiosity can therefore be operationalized as an intrinsic reward towards learning progress: the more a sequence is still teaching the infant something, the higher its instantaneous “value”, and the longer the MVT-driven agent should stay on it.

For each trial we therefore computed the learning progress that the trial offered to an ideal observer of the target locations, following^33^. The observer keeps track of how often the target has appeared at each of the four quadrants since the start of the current sequence, with a uniform Dirichlet prior:

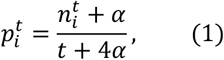

where 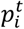 is the estimated probability that the next target lands in quadrant *i* after the first *t* trials of the sequence, which gets updated on every trial after stimulus presentation; 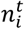 is the number of times quadrant *i* has been the target so far, and *α* is a pseudo-count expressing the strength of the prior. The learning progress contributed by trial *t* is then defined as the Kullback-Leibler divergence between the observer’s posterior after the trial and its posterior just before it:

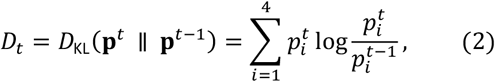

which captures how much the trial updated the observer’s beliefs about where the target tends to appear and is therefore taken as the amount of learning progress the infant could in principle extract from it^33^.

Given this definition of value, MVT can be implemented by maintaining a moving estimate of the local return on the current patch and comparing it to a moving estimate of the global return rate across the whole session. We track these two estimates with exponential moving averages (EMAs) of the per-trial learning progress signal *D*_*t*_ defined in Equation 2,

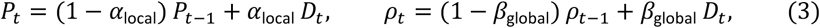

with *P*_0_ = *ρ*_0_ = 0. The local estimate *P*_*t*_ is updated at the fast rate *α*_local_, so it is dominated by the most recent trials and effectively reflects the current learning-progress reward value; the *global* estimate *ρ*_*t*_ is updated at the slower rate *β*_global_ and reflects the long-run average reward value. We parameterize the two rates by their *half-lives α*_hl_ = log2/*α*_local_ and *β*_hl_ = log2/*β*_global_, defined as the number of past trials after which the weight of an observation in the EMA has decayed to half its initial value.

The MVT signal at trial *t* is the difference between the local and global estimates, *X*_*t*_ = *P*_*t*_ − *ρ*_*t*_. Positive values indicate that the current sequence is yielding more learning progress than the running global average; negative values indicate that it has dropped below the global baseline, which is the regime in which an MVT-driven forager is predicted to leave.

#### Statistical analysis

We tested whether infants’ look-aways follow the MVT prediction by fitting a mixed-effects logistic regression of the look-away event on the MVT signal, pooling across the full sample with a random intercept by subject:

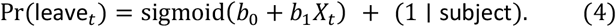

The free parameters of the framework are the EMA half-lives *α*_hl_ and *β*_hl_ that determine *P*_*t*_ and *ρ*_*t*_. We recovered them by grid search: for each integer pair *α*_hl_ ∈ {1, …, 10} trials and *β*_hl_ ∈ {1, …, 20} trials (constrained so that the global half-life is at least as long as the local one), we recomputed *P*_*t*_, *ρ*_*t*_, *X*_*t*_ and fitted a GLMER; the pair minimizing AIC was taken as the best fit. The MVT prediction is then assessed from the sign of the fixed effect *b*_1_: a negative coefficient indicates that the probability of leaving decreases with the MVT signal, consistent with the theory. The subject random intercept absorbs idiosyncratic baselines, so all infants, including those who happened to look away very few times, contribute to the shared fixed effect. We additionally fit a control mixed-effects model that augments the predictor set with the within-sequence and within-subject trial indices, so as to absorb pure time-on-task effects unrelated to the MVT signal.

The grid search selected a local half-life of 1 trial and a global half-life of 4 trials. At this optimum the probability of leaving fell steeply with the MVT signal: across the pooled sample (18,225 look-away decisions from 275 infants) the fixed effect of *X*_*t*_ was strongly negative (*β* = −0.80, *SE* = 0.03, *z* = −25.1, *p* < 0.001), the direction predicted by the theory: infants disengage once local learning progress falls below the running global baseline. The effect held in the control model that also absorbed within-sequence and within-subject trial position (*β*_*X*_ = −0.72, *SE* = 0.03, *z* = −20.7, *p* < 0.001), so it is not reducible to a simple decline with time on task (within-subject index *β* = 0.50, *SE* = 0.04, *z* = 12.8, *p* < 0.001; within-sequence index *β* = 0.11, *SE* = 0.03, *z* = 3.1, *p* = 0.002).

Finally, we compared our MVT model against two alternative theoretical accounts of disengagement that have been proposed in the literature ^33,34^. The first, a plain learning-progress account, predicts that infants leave when the current trial offers little learning progress *D*_*t*_, regardless of any baseline^33^. The second, a *Goldilocks* account^34^, predicts that infants engage with moderately surprising stimuli, predicting quadratic (inverted-U) effect of surprise *I*_*t*_ (the surprisal of the trial under the ideal observer’s pre-trial posterior). Both alternatives were fit on the same trials and the same random-intercept structure as the MVT model. Because the MVT model also selects (*α*_hl_, *β*_hl_) from a grid, we report a corrected AIC that adds 2 *⋅* 2 = 4 units to MVT’s raw AIC, treating the two grid-searched hyperparameters as additional model parameters. The MVT account clearly outperformed both alternatives, achieving the lowest AIC and BIC: it beat the plain learning-progress model by ΔAIC = 418 and the quadratic-surprise (“Goldilocks”) model by ΔAIC = 700, and its advantage persisted after penalizing the two grid-searched half-lives (ΔAIC ≥ 414, ΔBIC ≥ 398 against both).

### Recurrent Neural Networks

We trained RNNs with the MVT controller introduced above built directly into their training schedule, so that the order in which the models practice their tasks is decided online by the same kind of foraging rule we fitted to infants. The architecture and training pipeline are described in the Supplementary Information (Supplementary Text 1). Here, we focus on the foraging curriculum. We describe the MVT controller’s three ingredients in turn: the per-step measure of learning progress, the moving averages that summarize it, and the stay/leave decision rule.

#### Learning progress

At every gradient update on the current task *k*, the controller is given a scalar reward equal to the learning progress on that task, defined as the per-step reduction in the response-period cross-entropy:

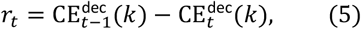

where 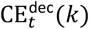 is the cross-entropy of the readout averaged across the response-period frames of the batch at update *t* on task *k*. Positive *r*_*t*_ means the loss has dropped (the network is improving on the task); negative *r*_*t*_ means it has risen. On the first visit to a task, 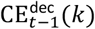 is undefined and *r*_*t*_ is set to zero. The raw reward is then passed through an asymmetric, prospect-theory-inspired value function before being handed to the controller,

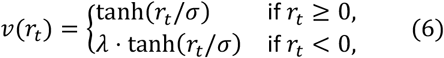

with *σ* = 0.1 controlling the width of the linear regime and *λ* = 2.25 amplifying the magnitude of losses relative to gains, consistent with prospect-theoretic accounts of decision-making under uncertainty^65^.

#### Local and global progress

Two exponential moving averages of the subjective reward *v*(*r*_*t*_) are then maintained at rates *α*_local_ and *β*_global_, exactly as in Equation 3: a local estimate *P*_*t*_ dominated by recent steps, and a global estimate *ρ*_*t*_ that tracks the long-run reward rate across the whole training run.

#### Decision rule

At every training step the controller commits to staying on the current task or leaving for a different one, drawing a stochastic decision from a soft version of the MVT rule:

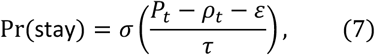

where *σ*(*⋅*) is the logistic function, *τ* is a decision temperature and *ε* is a leave margin. Regulating *τ* lets us continuously degrade the curiosity-driven preference encoded by the MVT signal: in the limit *τ* → 0 the rule becomes the deterministic MVT comparison (stay iff *P*_*t*_ > *ρ*_*t*_ + *ε*), so the controller is fully driven by learning progress; in the limit *τ* → ∞ the rule converges to a constant Pr(stay) = 0.5, so the controller picks tasks essentially at random: curiosity-driven exploration is replaced by random exploration. The margin *ε* shifts the threshold for staying: negative values make the controller stickier (it tolerates some plateau before leaving), positive values make it more eager to switch.

#### Travel cost

When the controller commits to leaving, the network enters a *travel* phase of *n*_travel_ steps during which no gradient update is performed but the global baseline *ρ*_*t*_ continues to update under *r*_*t*_ = 0. After travel, the next task is drawn uniformly at random from the remaining tasks, excluding the one just left. Because *ρ*_*t*_ decays during travel, longer travels leave the agent with a lower estimate of the global return rate; as a consequence, when starting the next task, more modest local progress will still be desirable compared to the low global average, and the controller will tend to stay longer in the patch. This is the classical MVT result that, when travel between patches is expensive, optimal dwell time within each patch increases. Travel steps do not consume the total budget of training steps.

All parameters specified in the previous sections are held fixed at the values stated there, with two exceptions: the choice of weight regularizer and the five hyperparameters of the MVT controller. These are the parameters we deliberately leave free across the sweep, either because their best values cannot be determined a priori, and/or because their effect on the trained models is precisely what we want to characterize. A complete summary of all the free parameters and their values is available in the Supplementary Information (Supplementary Text 2).

### Analysis

We assess the trained networks along three axes: network structure, synaptic density, and compositional generalization.

#### Network structure

In terms of network structure, we compared the topology of each trained RNN’s recurrent connectivity to the topology of the human structural connectome. The preprocessing steps to obtain weighted adjacency matrices from RNNs and human connectome data are described in the Supplementary Information (Supplementary Text 3). We computed three graph-theoretic measures^66^: modularity, global efficiency, and rich-club coefficient. Together, they summarize complementary aspects of network organization – segregation, integration, and hierarchical structure – providing a compact, three-dimensional fingerprint of a connectome’s topology. More details on the topological measures are available in the Supplementary Information (Supplementary Text 4).

To quantify how brain-like an RNN’s topology is, we compared its fingerprint to the average human fingerprint (defined as the mean fingerprint across the individual HCP-YA participants). First, we used Mahalanobis distance to obtain a distance score between each RNN and the human average (see also Supplementary Text 4). Then, we obtained a similarity score by linearly rescaling distance scores to a similarity *s* ∈ [0,1] (*s* = 1 for the most human-like network, *s* = 0 for the least). This score is the variable used in the regression and enrichment analyses below.

We asked which of the manipulated components made a network’s connectome more brain-like, using the network-similarity score *s* as the outcome. The predictors were the three components varied across the sweep: the foraging temperature *τ*, the regularization strength, and the regularization type (distance-based vs. L1). We fitted the full three-way factorial linear model and assessed term significance with Type-II analysis of variance, which is appropriate for the mildly unbalanced design. Because the weakest regularization level existed only for L1, it was excluded so that the design was fully crossed (seven temperatures × three strengths × two regularization types). We then performed Tukey-corrected post-hoc tests on the estimated marginal means. The model explained 86% of the variance in network similarity (*F*_41, 2338_ = 360.3, *p* < 0.0001). Crucially, the three-way interaction between the three components was significant (*F*_12, 2338_ = 5.3, *p* < 0.0001), indicating that the effect of temperature depended jointly on the type and strength of regularization.

Post-hoc tests on the estimated marginal means located the optimum. Within the distance-based, strongest-regularization regime, predicted similarity peaked at *τ* = 0.003 (estimated marginal *s* = 0.81, 95% CI [0.79,0.82]), the most brain-like cell of the entire design. In this regime the curiosity-driven temperatures were the most human-like: *τ* = 0.003 exceeded the random regime *τ* = 1.0 by Δ*s* = 0.06 (Tukey *p* < 0.0001) and exceeded every higher temperature *τ* ≥ 0.01 (all Tukey *p* < 0.0001), while the lowest temperature *τ* = 0.0001 was statistically indistinguishable from the peak (*s* = 0.80, Tukey *p* ≈ 1). This temperature profile was specific to the strongly, distance-regularized networks: under L1 regularization the predicted similarity across the same temperatures spanned only *s* ≈ 0.40–0.42 and was essentially flat, consistent with the significant three-way interaction.

To check that the result does not depend on the four remaining hyperparameters of the MVT controller (the local progress rate *α*_local_, the global baseline rate *β*_global_, the travel cost *n*_travel_, and the leave margin *ε*), we re-fitted the model with each of the four hyperparameters in turn fully crossed with the three components. The three-way interaction remained significant in every case (all *p* < 0.0001), and a regime with curiosity-driven temperature and strong, distance-based regularization remained the predicted optimum across all 64 combinations of the four hyperparameters.

Next, we asked which parameter values are over-represented among the networks that actually reached the most brain-like topology (enrichment analysis). From the full functional pool, we took the 100 networks with the highest network-similarity score *s* (equivalently, the smallest Mahalanobis distance to the average human fingerprint, i.e., the mean across the individual HCP-YA networks). For each swept parameter and each of its values we computed an enrichment factor, the proportion of the top-100 networks taking that value divided by its proportion in the full pool, so that a factor above one indicates over-representation among the most brain-like networks. For each parameter we tested whether the distribution of values among the top-100 differed from the pool with a *χ*^2^ test whose *p*-value was obtained by Monte-Carlo simulation (*B* = 10,000 resamples), which is exact for the small expected counts in some cells. The composition of the 100 most brain-like networks differed strongly from the pool on all three components: regularization type (*χ*^2^ = 145.5, *p* < 0.0001), regularization strength (*χ*^2^ = 211.8, *p* < 0.0001) and temperature (*χ*^2^ = 76.6, *p* < 0.0001). The over-represented values were consistently distance-based regularization, the strongest regularization, and the curiosity-driven temperatures, with L1, weak regularization and the random regime correspondingly depleted.

#### Synaptic density

We aimed at investigating whether RNNs that are more human-like in terms of foraging and biophysical constraints also display more human-like developmental trajectories of synaptic density and cognitive performance. For RNNs, we tracked the density of recurrent weights across training; for humans, the density of synapses across the lifespan. Both quantities measure how densely a system is wired at a given moment. Details on how these quantities were extracted are available in the Supplementary Information (Supplementary Text 5).

The analysis has two goals. The first is to test whether fitted age differs between the 100 networks most similar to the human connectome in topology (smallest network Mahalanobis distance; see the network analysis above) and the rest of the RNNs (*N* = 2,704): we expect the most similar networks to have more plausible fitted ages, capturing more of the lifespan trajectory. To test this we compared the distribution of fitted ages in the two groups with a *χ*^2^ test, with the *p*-value obtained by Monte-Carlo simulation (*B* = 10,000 resamples) given the small expected counts in some age bins. The two groups had markedly different fitted-age distributions (*χ*^2^ = 88.7, *p* < 0.0001). The most human-like networks were assigned substantially older ages (median fitted age 32 vs 9 years): their weight trajectories tracked a fuller developmental arc, an extended rise-and-prune course, whereas the rest mostly matched only a brief early window.

The second goal is to relate the inferred developmental trajectory to task performance. To test this, within the 100 most human-like networks we modeled end-of-training accuracy as a smooth, potentially non-linear function of fitted age with a generalized additive model. We found a strong, non-linear effect (*edf* = 2.63, *F* = 15.9, *p* < 0.001, 35.5% of deviance explained): accuracy rose with fitted age to a peak at ≈ 20 years and declined thereafter, recovering the characteristic rise and gradual decline of cognitive performance across the human lifespan.

#### Compositional generalization

Compositional generalization consists in the capacity to handle novel arrangements of familiar parts^46,49^. Testing this ability cannot rely on zero-shot performance, whereby an agent is tested directly on new tasks with no further training. The reason is that a network cued for a task it has never seen, much like a person given a new task with no instructions, needs a short period of trial and error simply to establish what is being asked. We therefore allow a brief adaptation phase, but constrain it so that the network cannot learn the held-out task from scratch: it can succeed only by recruiting recurrent dynamics it already has. To this end every trained weight is frozen and a single new rule-input channel is exposed for adaptation. This way, a held-out task can be solved only by routing the new input into existing recurrent computations, that is, by recombining primitives the network already possesses.

Compositional reuse is therefore indexed by the *speed* of adaptation rather than by the accuracy eventually reached. To make that speed interpretable, we applied the same procedure to tasks the network had already been trained on, again withholding any cue that they were familiar: if a network’s internal dynamics for a known task remain intact and reachable, it should re-solve that task almost immediately. A held-out recombination solved as quickly as such a familiar task thus reflects genuine reuse of existing dynamics, whereas one solved slowly indicates the solution is being rebuilt through the new channel. Compositional generalization is demonstrated only when a network solves both a novel recombination and its familiar reference rapidly: the bottom-left corner of the solve-time plane (Fig. 4G-H). All details on how the held-out tasks were selected and the test was carried out are available in the Supplementary Information (Supplementary Text 6).

To quantify the speed of resolution, measured how many steps were necessary to reach 90% accuracy on training (*t*_train_) and held-out tasks (*t*_held_). Each pair of measures (*t*_train_, *t*_held_) was summarized by the geometric mean of its two solve times, 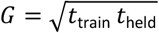 (Figure 4G-H). In this parameterization a positive coefficient denotes a longer (slower) solve time.

To disentangle the contributions of the two human-like components (curiosity-driven exploration and brain-like spatial constraints) we compared four groups of trained networks. The human-like group consisted of the 100 networks most similar to human topology; Among those, we retained the ones that had learned the task battery best (fraction of training tasks solved ≥ 0.9). These networks lay in the curiosity-driven (temperature *τ* < 0.01), strongly regularized regime. For each retained network, three matched control networks were then drawn from the same trained population, sharing all hyperparameters with their human-like counterpart except for one or both of the two human-like components: the Temperature Control is the network whose foraging temperature is instead *τ* = 1.0 (an essentially random task-selection regime); the Regularization Control is the network whose regularization strength is instead the minimum value available; and the Both Control is the network with both substitutions. The four groups therefore cross temperature (low vs random) with regularization strength (high vs minimal). The cohorts contained 25 to 30 networks each, for a total of 110 networks tested.

We fitted a log-normal accelerated-failure-time model on our variable of interest *G*. The model included temperature (curiosity, *τ* < 0.01, vs random, *τ* = 1), regularization (strong vs minimal), and their interaction with the type of ability tested. Finally, differences between cohorts could be confounded by how well each network had learned the relevant abilities by the end of training. We therefore added the network’s end-of-training accuracy on the base abilities probed by the test. Curiosity-driven exploration (low temperature) led to faster resolution in all three tests (*distractor* added to a decision, *β* = −0.50, *SE* = 0.058, *t* = −8.60, *p* < 0.001; *anti-mapping* added to a match, *β* = −0.28, *SE* = 0.072, *t* = −3.97, *p* < 0.001; *distractor* and *anti-mapping* added to a match, *β* = −0.30, *SE* = 0.072, *t* = −4.18, *p* < 0.001). In addition, strong regularization solved faster than minimal regularization (*β* = −0.41, *SE* = 0.039, *t* = −10.66, *p* < 0.001).

## Acknowledgements

Connectomics data were analysed using the US National Science Foundation, ACCESS program, resource allocation (TG-CIS200026) at Extreme Science and Engineering Discovery Environment (XSEDE) resources (Towns, J. et al. Computing in science & engineering 16, 62-74 2014) by Fang-Cheng Yeh at the University of Pittsburgh.

## Author contributions

F.P.: Conceptualization, Methodology, Investigation, Formal analysis, Data Curation, Writing – Original draft, Visualisation, Funding acquisition; K.F.: Conceptualization, Formal analysis, Writing – Review & Editing; A.M.: Data Curation, Writing – Review & Editing; C.K: Investigation, Data Curation, Writing – Review & Editing; C.G.W.: Resources, Writing – Review & Editing; D.A.: Conceptualization, Methodology, Resources, Writing – Review & Editing, Supervision, Funding acquisition.

## Data and materials availability

Data, code, and materials are available on Github: https://github.com/FrancescPoli/multitask-curiosity-release

