## Supplementary Information for "Curiosity shapes brain-like architectures and functions"

This document includes the following content:


Supplementary Text 1: Recurrent Neural Network Models 2

Supplementary Text 2: Free parameters 8

Supplementary Text 3: Network data processing 9

Supplementary Text 4: Topological measures 11

Supplementary Text 5: Synaptic density data and preprocessing 13

Supplementary Text 6: Implementation of the compositional generalization test 14

References 15

### Supplementary Text 1: Recurrent Neural Network Models

Our network builds on the multi-task recurrent architecture introduced by Yang and colleagues[1] and adopts the unified categorical readout used by Khona and colleagues[2], while differing from both in the choice of training objective, regularization, and training curriculum (described in the following sections). It consists of a single hidden layer of $N = 100$ leaky-integrator units with rectified linear (ReLU) activation, which receive a structured task input at every timestep and project, through a linear readout followed by a softmax, onto a discrete action space. At each timestep the input vector and the recurrent state are concatenated and enter the hidden layer through a single shared weight matrix, so that sensory and task-identifying channels are integrated with the recurrent state through the same set of trainable connections.

#### Recurrent dynamics

Hidden activity $\mathbf{h}_t \in \mathbb{R}^N$ evolves according to an Euler-discretized leaky-integrator equation:

$$\mathbf{h}_{t+1} = (1-\alpha)\,\mathbf{h}_t \;+\; \alpha\, f(\mathbf{W}\,[\,\mathbf{x}_t;\, \mathbf{h}_t\,] + \mathbf{b} + \boldsymbol{\xi}_t^{\text{rec}}), \qquad (1)$$

where $\mathbf{x}_t$ is the input vector at time $t$, $[\,\cdot\,;\cdot\,]$ denotes concatenation, $\mathbf{W}$ collects the fused input-to-hidden and recurrent weights, $\mathbf{b}$ is a bias vector, and $f$ is the elementwise ReLU nonlinearity. The leak coefficient is set by the ratio of the discretization step to the membrane time constant, $\alpha = \Delta t/\tau_m$. We use $\Delta t = 20$ ms and $\tau_m = 100$ ms, giving $\alpha = 0.2$. The hidden state is reset to $\mathbf{h}_0 = \mathbf{0}$ at the start of every trial.

Two independent sources of zero-mean Gaussian noise are injected during training. The first, $\boldsymbol{\xi}_t^{\text{rec}}$ in Equation 1, acts directly on the pre-activation of each recurrent unit and is intended to capture the intrinsic stochasticity of biological neurons. The second is added to the input vector $\mathbf{x}_t$ before it enters the recurrent map and represents perceptual uncertainty at the sensory boundary, mimicking unreliable observations and inter-trial variability in the signals the network is exposed to. The two noise sources have standard deviations $\sigma_{\text{rec}}\sqrt{2/\alpha}$ and $\sigma_x\sqrt{2/\alpha}$, with $\sigma_{\text{rec}} = 0.05$ and $\sigma_x = 0.01$. The $\sqrt{2/\alpha}$ scaling ensures that the effective noise variance at the membrane level is preserved across choices of $\Delta t$ [1]. Both noise sources are disabled at evaluation time, so any stochasticity in the model's behavior at test reflects only the noise it experienced during training.

#### Input

The input vector $\mathbf{x}_t$ presented to the network at each timestep is the concatenation of three functional blocks: a fixation cue, sensory channels carrying the current stimulus, and a static task-identifying signal. The total input has 133 dimensions (1 fixation channel, 32 sensory channels and 100 task-identifying channels), and all three blocks enter the hidden layer through the same shared weight matrix introduced above.

**Fixation cue.** A single binary-valued channel encodes a fixation cue. It is set to 1 during the periods of the trial in which the network is required to maintain fixation – typically the fixation, stimulus and (when present) delay periods – and switches to 0 once the network is allowed to commit a motor response. The exact mapping between trial periods and the cue's value differs slightly across task variants, as described later.

**Sensory channels.** The current stimulus is encoded across two rings of 16 directional channels each, for a total of 32 sensory channels. Each of the 16 channels is assigned a direction it responds most strongly to, with the preferred directions equally spaced around the circle (see Fig. 1A). A channel fires not only at its preferred direction but with declining strength as the stimulus moves away from it. A single stimulus therefore activates the closest channel plus a few of its neighbors, with smoothly graded magnitudes.

Concretely, for a stimulus at orientation $\theta$ the activity of the $k$-th channel in a ring is

$$s_k = 0.8 \cdot \exp\left(-\frac{1}{2}\left(\frac{d(\theta_k, \theta)}{\pi/8}\right)^2\right), \qquad (2)$$

where $\theta_k = 2\pi k/16$ for $k = 0, \dots, 15$ indicates the 16 equally spaced positions on the ring, $\theta_k$ indicates the preferred direction of channel $k$, and the declining strength of response is captured by the Gaussian envelope with width $\pi/8$, which matches one inter-channel spacing. The angular distance $d(\theta_k, \theta)$ between the channel's preferred direction and the stimulus is the periodic distance on the circle.

**Task-identifying channels.** In the original multi-task RNN[1], the task to be executed on each trial was signaled to the network through a one-hot rule input: a vector with as many channels as tasks, in which exactly one channel was set to $1$ and all others to $0$. Later work[2] showed that this scheme biases the network toward a degenerate solution in which the multi-task battery is solved largely through *autapses*, i.e., strong recurrent self-connections: with a private input dimension per task, the network can simply learn task-specific units that are driven by their dedicated rule channel and need only maintain their own activity to produce the correct output, with little genuine inter-unit recurrent processing. This bypasses the computational role of the recurrent layer and prevents the emergence of a shared representational substrate across tasks.

We instead use a dense, distributed rule code, designed so that the task-identifying input is necessarily ambiguous. Each task is assigned a vector that is drawn at random in a low-dimensional latent space (four dimensions) and then projected up to the $100$ input channels through a fixed random map. Because $20$ vectors cannot be packed orthogonally into a space of only $4$ dimensions, the task vectors necessarily overlap, with an expected absolute cosine similarity of roughly $1/\sqrt{4} = 0.5$ between any two of them. Which tasks happen to overlap more or less is set entirely by the random draw. The random rule geometry therefore carries no information about task structure: that structure must be discovered by the network from the loss alone.

When the network receives the rule vector for a given task, the input drive partially overlaps with the drive that would be produced by the rule vectors of several other, unrelated tasks, and the corresponding subsets of hidden units are co-activated even though only one task is the actual target. Hence, the network must use its recurrent connectivity to perform pattern separation, leveraging lateral inhibition and other forms of inter-unit coupling to suppress the interfering task signals and amplify the target one. The autaptic solution becomes prohibitively expensive, and the network is pushed toward genuinely recurrent, distributed computation in which units are reused across tasks.

Concretely, the rule vector $\mathbf{r}^{(k)}$ for task $k$ is constructed in two stages. First, a 4-dimensional latent $\mathbf{z}^{(k)}$ is drawn from a standard Gaussian and renormalized to unit length. Second, it is projected to $100$ dimensions through a fixed random Gaussian matrix $\mathbf{P} \in \mathbb{R}^{4\times 100}$ (each entry sampled iid from $\mathcal{N}(0,1)$), and the resulting 100-dimensional vector is again renormalized to unit length:

$$\mathbf{r}^{(k)} = \frac{\mathbf{P}^\top \mathbf{z}^{(k)}}{\| \mathbf{P}^\top \mathbf{z}^{(k)} \|}. \qquad (3)$$

Each task is associated with a single rule vector. Both $\mathbf{P}$ and the set of latents $\{\mathbf{z}^{(k)}\}$ are drawn once at the start of training using a fixed seed, held constant thereafter, and not optimized during training.

### Initialization

All trainable parameters of the recurrent layer are initialized once, under the same random seed that governs the rule-vector projection. Input weights (rows of $\mathbf{W}$ in Equation 1 acting on $\mathbf{x}_t$) are drawn independently from a zero-mean Gaussian with variance $1/n_{\text{input}}$, so that the variance of the total input drive to each unit is of order one. Recurrent weights (rows acting on $\mathbf{h}_t$) are initialized as a random orthogonal matrix rescaled so that the spectral radius of the recurrent map is 0.5, keeping the network in a sub-critical regime in which spontaneous activity decays in the absence of input. Biases are initialized to zero.

### Readout

At each timestep the hidden state $\mathbf{h}_t$ is mapped, through a linear projection followed by a softmax, to a probability distribution over a discrete action space of 17 classes: one *fixation* class, corresponding to the network withholding a motor response, and 16 *direction* classes, one for each of the ring directions used in the sensory input. Concretely,

$$\mathbf{l}_t = \mathbf{W}_{\text{out}}\,\mathbf{h}_t + \mathbf{b}_{\text{out}}, \qquad \mathbf{p}_t = \text{softmax}(\mathbf{l}_t), \qquad (4)$$

with $\mathbf{W}_{\text{out}} \in \mathbb{R}^{17\times N}$ initialized by Xavier-uniform sampling and $\mathbf{b}_{\text{out}} \in \mathbb{R}^{17}$ initialized to zero. The readout is applied at every timestep of the trial, so the network produces a class distribution throughout the trial and can be supervised against the target action at every step.

### Loss function

Training minimizes a composite loss combining a temporally weighted supervised term on the readout with a single weight-regularization term, the latter chosen across runs to lie in one of two regimes.

**Supervised term.** At every timestep, the categorical distribution $\mathbf{p}_t$ produced by the readout is supervised against the target action $y_t \in \{0,1,\ldots,16\}$ provided by the task generator. We follow the temporal-weighting scheme of Yang et al. (2019): the per-trial loss is

$$\mathcal{L}_{\text{CE}} = \frac{1}{|\mathcal{V}|}\sum_{t\in\mathcal{V}} w_t\ \text{CE}(\mathbf{p}_t, y_t), \qquad w_t = \begin{cases} 5 & \text{if } y_t \in \{1,\ldots,16\} \\ 1 & \text{if } y_t = 0 \end{cases} \qquad (5)$$

where $\text{CE}(\mathbf{p}_t, y_t) = -\log p_{t,y_t}$ is the categorical cross-entropy at frame $t$ and $\mathcal{V}$ is the set of frames at which a target action is defined (padding frames inserted to align trial lengths within a batch are excluded). Response frames, those at which the target is one of the 16 direction classes, therefore contribute five times as much to the loss as fixation frames; this prevents the loss from being dominated by the typically long fixation periods, over which the readout target is the same class at every frame.

**Regularization.** A single weight-regularization term $\mathcal{R}$ is added to the supervised loss, so that the total optimized in a given run is $\mathcal{L} = \mathcal{L}_{\text{CE}} + \mathcal{R}$. We consider two regimes:

*L1.* An $\ell_1$ penalty applied uniformly across all trainable weights:

$$\mathcal{R}_{\text{L1}} = \lambda_{\text{L1}} \sum_{\theta\in\Theta_{\text{w}}} |\,\theta|, \qquad (6)$$

where $\Theta_w$ collects the entries of $\mathbf{W}$ (both input and recurrent blocks) and of $\mathbf{W}_{out}$; biases, being one-dimensional, are excluded. This regime applies a uniform pressure toward sparsity throughout the network.

*Distance-weighted penalty.* An $\ell_1$ penalty applied only to the recurrent weights, with each weight multiplied by a fixed cost that grows with the spatial distance between the two units it connects:

$$\mathcal{R}_{\text{dist}} = \lambda_{\text{dist}} \sum_{i,j=1}^{N} |W_{ij}^{\text{rec}}| D_{ij}^2, \qquad (7)$$

where $W_{ij}^{\text{rec}}$ is the recurrent weight from unit $j$ to unit $i$. To assign distances to the units, we identify the $N = 100$ hidden units with the $100$ cortical parcels of the Schaefer-100 atlas[3] and take $D_{ij}$ to be the Euclidean distance between the centroids of parcels $i$ and $j$, normalized so that $\max_{i,j} D_{ij} = 1$. The squared exponent makes the cost grow rapidly with distance, encouraging the recurrent connectivity to remain spatially local in a way reminiscent of the distance-dependent wiring cost observed in cortex[4].

### Optimization

The composite loss is minimized by stochastic gradient descent with Adam[5] at a learning rate of $1 \times 10^{-3}$. At each step, a batch of $16$ independent trials is drawn from the task currently selected by the curriculum (described below), and a single gradient update is performed. To prevent rare large gradients from destabilizing the recurrent dynamics, gradients are clipped element-wise to $[-1, +1]$ before the optimizer step. Training consisted of a total of $6 \times 10^5$ gradient updates.

Cognitive Tasks

The network is trained on a battery of $20$ cognitive tasks adapted from an existing multi-task suite[1]. All $20$ tasks share the same input format, action space and temporal structure, but differ in the rules that map sensory input to the correct response. They are organized into three behavioral families – *reaching*, *perceptual decision-making* and *delayed match-to-sample* – and parameterized by a small set of modifiers, or primitives (anti, ctx, dly, rt, multi, cat, nms), which combine to generate the variants within each family.

**Trial structure.** Every trial unfolds over four sequential periods, presented to the network as consecutive timesteps of $\Delta t = 20$ ms:

1. **Fixation period**: $300$-$700$ ms, drawn uniformly per trial. The fixation cue is set to $1$, both sensory rings are silent, and the correct action is the *fixation* class.
2. **Stimulus period**: $500$-$1500$ ms uniform (split into separate *sample* and *test* sub-periods of $200$-$600$ ms each in match and delayed-decision tasks). One or two Gaussian bumps are presented on the sensory rings, as required by the task.
3. **Optional delay period**: $500$ ms in the reaching family; $200$ - $1600$ ms (drawn from $\{200, 400, 800, 1600\}$) in match and delayed-decision tasks. The fixation cue remains $1$, both rings are silent, and the correct action is still *fixation*. This period is absent in tasks without an explicit delay modifier.
4. **Response period**: $500$ ms. The fixation cue switches to $0$ and the correct action becomes one of the $16$ direction classes, determined by the task rule below.

Period durations are resampled independently on every trial. Each batch element is a fresh, independent trial starting from a zero hidden state, so the network never carries state from one trial to the next.

**Reaching family ($10$ tasks).** A single Gaussian-bump stimulus is presented at orientation $\theta$, and the network must respond at the matching direction during the response period. In tasks without the context modifier, the stimulus is placed on either ring 1 or ring 2 with equal probability per trial, so the network must learn to extract direction independently of which ring carries the signal. Four binary modifiers generate the ten reaching variants:

- **Anti-Mapping (A)**: the correct response is rotated by $\pi$ (the network must reach in the direction opposite to the stimulus).
- **Distractor (D)**: two stimuli are presented simultaneously, one on each ring; the network must respond to the stimulus on ring 1 (the target) and ignore the stimulus on ring 2 (the distractor).
- **Delay Interval (I)**: a delay period is inserted between stimulus offset and response; the network must hold the stimulus direction in working memory across a stimulus-free interval before committing.
- **Reaction Time (R)**: the fixation cue drops at stimulus onset and the response is read out continuously from the stimulus period onward, rather than waiting for a fixed response window.

The ten reaching tasks correspond to the combinations actually used in our battery: go, Rgo, Igo, Ago, Dgo, IAgo, IDgo, ADgo, IADgo, RADgo.

**Perceptual decision-making family ($6$ tasks).** Two Gaussian-bump stimuli are presented at opposite orientations with different *coherences* (effective contrasts) $c_1$ and $c_2$, drawn so that $c_1 + c_2 = 1$ and $|c_1 - c_2| \in \{0.08, 0.16, 0.32\}$ per trial; smaller differences make the trial more ambiguous. The "winner" is the more coherent of the two stimuli, and the correct response is its direction. Stimulus channels are additionally corrupted by zero-mean Gaussian noise during the stimulus period, so coherence sets the per-frame signal-to-noise ratio that the network must integrate over time. The two stimuli are presented on ring 1 only (dm1) or ring 2 only (dm2). Three modifiers generate the six decision-making variants:

- **Anti-Mapping (A)**: the target is rotated by $\pi$, as in the reaching family.
- **Delay Interval (I)**: a delay separates a brief *sample* presentation from a *test* presentation, as in the reaching family.
- **Multisensory (M)**: the stimuli are presented redundantly on both rings; the network must combine evidence across modalities.

The six tasks are: dm1, dm2, Adm, Ddm1, Mdm, Idm1, ADdm1.

**Delayed match-to-sample family ($4$ tasks)**. A *sample* stimulus at orientation $\theta_s$ is presented, then a delay, then a *test* stimulus at orientation $\theta_t$. The network must commit to the *test* direction if a task-specific match condition is satisfied, and otherwise maintain fixation throughout the response period. In its basic form, the trial can either be **Match-to-Sample (ms)** or **Non-Match-to-Sample (nms)**. Under ms the network responds at $\theta_t$ when $\theta_t = \theta_s$; under nms the network responds at $\theta_t$ when $\theta_t \neq \theta_s$. The two types are drawn with equal probability per trial. Two binary modifiers generate the four variants:

- **Delay Interval (I)**: In match-to-sample tasks, a delay always separates the sample stimulus from the test stimulus.

- **Categorical Rule (C)**: the match condition is relaxed from exact identity to same-hemisphere: $\theta_s$ and $\theta_t$ are considered to match if they both fall in the upper half-circle $(0, \pi]$ or both in the lower half-circle $(\pi, 2\pi]$. This requires the network to learn an abstract categorization rule rather than an exact sample–test identity rule.

All four match tasks include a delay between sample and test: Ims, Inms, CIms, CInms.

### Supplementary Text 2: Free parameters

Five hyperparameters were systematically manipulated. First, we varied both the *type* and the *coefficient* of the weight regularizer. The type takes one of two values: an $\ell_1$ penalty or a distance-weighted $\ell_1$ penalty. The two types are mutually exclusive: each run uses either an $\ell_1$ penalty with $\lambda_{\text{L1}} \in \{10^{-6}, 10^{-5}, 10^{-4}, 5 \cdot 10^{-4}\}$, or a distance penalty with $\lambda_{\text{dist}} \in \{10^{-5}, 10^{-4}, 5 \cdot 10^{-4}\}$.

In addition, five hyperparameters of the MVT controller are varied:

- **Decision temperature** $\tau \in \{10^{-4}, 10^{-3}, 3 \cdot 10^{-3}, 5 \cdot 10^{-3}, 10^{-2}, 10^{-1}, 1.0\}$.
- **Local progress rate** $\alpha_{\text{local}} \in \{10^{-2}, 3 \cdot 10^{-2}, 10^{-1}, 2 \cdot 10^{-1}\}$.
- **Global baseline rate** $\beta_{\text{global}} \in \{10^{-4}, 10^{-3}, 3 \cdot 10^{-3}, 10^{-2}\}$.
- **Travel cost** $n_{\text{travel}} \in \{50, 500\}$.
- **Leave margin** $\varepsilon \in \{0, -0.01\}$.

The temperature $\tau$ is varied across four orders of magnitude to map the entire deterministic-to-random regime of the soft MVT rule. The remaining hyperparameters were varied because no previous work has trained RNNs using MVT, so any a priori choice would have been arbitrary; sweeping each of them lets us characterize how MVT-driven curricula shape both the training trajectories and the properties of the resulting networks. The rates $\alpha_{\text{local}}$ and $\beta_{\text{global}}$ jointly set the timescale separation between local and global reward estimates, which determines how much past performance is integrated before a switch can be triggered. The travel cost $n_{\text{travel}}$ is varied between a cheap and an expensive switching regime, leading to curricula with different average dwell times per task. The margin $\varepsilon$ takes either zero (neutral) or a small negative value (mildly sticky).

The two regularization branches yield $4 \times 4 \times 4 \times 2 \times 7 \times 2 = 1\,792$ configurations under $\ell_1$ and $3 \times 4 \times 4 \times 2 \times 7 \times 2 = 1\,344$ configurations under distance, for a total of $3\,136$ trained models. Each model is trained from a single random seed.

### Supplementary Text 3: Network data processing

**RNN preprocessing.** For each trained network we take a single connectivity matrix and treat it as the RNN's "connectome". The relevant weights are the recurrent block of the fused weight matrix $\mathbf{W}$ introduced in Equation 1, namely the $N \times N$ sub-matrix $W^{\text{rec}}$ acting on the hidden state $\mathbf{h}_t$. We discard the input-to-hidden block, since it carries sensory and rule signals rather than inter-unit recurrent connectivity, and we discard the readout weights $\mathbf{W}_{\text{out}}$ for the same reason.

We then construct a non-negative weighted adjacency matrix $A^{\text{RNN}}$ by taking the element-wise absolute value of the recurrent block and symmetrizing it,

$$A_{ij}^{\text{RNN}} = 1/2\,(|W_{ij}^{\text{rec}}| + |W_{ji}^{\text{rec}}|), \qquad A_{ii}^{\text{RNN}} = 0, \qquad (8)$$

with the diagonal set to zero so that autaptic self-connections do not contribute to the topology, as it is standard practice in human brain connectomics. Discarding the sign means that we treat excitatory and inhibitory recurrent connections as topologically equivalent edges, also matching the convention adopted on the human side, where streamline counts are sign-less. Given that the recurrent weight matrix was directed ($W_{ij}^{\text{rec}} \neq W_{ji}^{\text{rec}}$ in general), we averaged the two directions yielding a symmetric matrix that can be treated as an undirected weighted graph, exactly like the human connectome.

The procedure yields one $100 \times 100$ adjacency matrix per trained run. The analysis pool covers all sweep configurations that learned the task battery to a basic functional level, operationalized as a fraction of solved tasks of at least $0.3$. This threshold is deliberately loose: it removes networks that failed to train altogether while keeping the bulk of the sweep, including networks that learned only a subset of the tasks.

**Human imaging data preprocessing.** Human connectomes were generated from the Human Connectome Project Young Adult (HCPya) dataset [6]. This sample (N = 1065; age range 22-37 years; 54% female) is a large sample of healthy adults. High-quality multishell diffusion-weighted MRI (dMRI; b-values = 1000 s/mm$^2$, 2000 s/mm$^2$, 3000 s/mm$^2$; 90 sampling directions each) was collected by the Washington University-University of Minnesota Consortium [7]. This data was accessed in a semi-reconstructed format from DSI Studio's Fiber Data Hub [8]. The DSI Studio team took the minimally preprocessed dMRI data and reconstructed it in MNI space using q-space diffeomorphic reconstruction [9]. We utilized the GQI reconstructed FIB files.

Tractography was conducted using the 100-region Schaefer atlas [3] using DSI Studio's whole-brain deterministic tractography [10]. Tracking parameters were: 35 degree turning angle, 5 million maximum fiber count, 1.0mm step size, 30mm minimum, and 250mm maximum fiber length. The streamlines were counted using count-pass connectivity, which identifies regions as connection if the streamline passes through that region. Fiber tracking yield 100 x 100 adjacency matrices per participant.

**Joint preprocessing.** Across the HCP-YA cohort, the density of the Schaefer-100 connectomes ranged from $0.138$ to $0.200$ (mean $0.169$, SD $0.010$). We thresholded every network – both individual HCP-YA connectomes and every RNN matrix – to the common target density $\rho^\star = 0.135$, just below the lower bound of this empirical distribution, so that the threshold could be applied uniformly to all human and RNN networks without artificially sparsifying any subject's connectome. Thresholding was performed by ranking the off-diagonal upper-triangular entries by absolute weight and retaining the top $\lfloor \rho^\star \cdot N(N-1)/2 \rfloor = 668$ edges in each network, setting the rest to zero. After thresholding, the surviving edge weights of every network were divided by their mean, so that each connectome is expressed on a common, unit-free weight scale. This mean-normalization makes the weighted topological measures scale-invariant, so that differences between RNN and human networks reflect the relative organization of their connections rather than the absolute magnitude of the weights (which differ arbitrarily between recurrent weights and streamline counts).

### Supplementary Text 4: Topological measures

On each thresholded network we computed three graph-theoretic measures, chosen because they are largely independent of each other and are widely used to characterize biological brain connectomes [11]: modularity, global efficiency, and rich-club coefficient. Together, they summarize complementary aspects of network organization – segregation, integration, and hierarchical structure – providing a compact, three-dimensional fingerprint of a connectome's topology.

*Modularity* [12] measures the extent to which the network decomposes into densely connected sub-communities with comparatively few connections between them, and is therefore an index of *segregation*. We partition the thresholded weighted graph into communities using the Leiden algorithm [13], which improves on the widely used Louvain algorithm [14] by guaranteeing well-connected communities, and report the modularity $Q$ of the recovered partition:

$$Q = \frac{1}{2m} \sum_{i,j} \left[ A_{ij} - \frac{k_i k_j}{2m} \right] \delta(c_i, c_j), \qquad (9)$$

where $A_{ij}$ is the (thresholded, weighted) adjacency matrix, $k_i = \sum_j A_{ij}$ is the strength of node $i$, $m = 1/2 \sum_{ij} A_{ij}$ is the total network strength, $c_i$ is the community assigned to node $i$, and $\delta$ is the Kronecker delta. Higher $Q$ indicates stronger community structure.

*Global efficiency* [15] measures the typical ease of communication between any two nodes, treating each edge as a conductance through which signals propagate, and is therefore an index of *integration*. To compute it on a weighted graph, edge weights are first inverted to distances, $d_{ij} = 1/A_{ij}$, and the shortest-path distance $L_{ij}$ between every pair of nodes is computed under the resulting metric. The global efficiency is then the average inverse shortest-path distance,

$$E = \frac{1}{N(N-1)} \sum_{i \neq j} \frac{1}{L_{ij}}, \qquad (10)$$

so that strongly connected pairs of nodes – connected either directly or through short, high-weight paths – contribute more, and pairs that are only reachable through long or weak detours contribute less.

*Rich-club organization* [16,17] measures the extent to which high-degree nodes are preferentially connected to one another, forming a densely interconnected hub backbone, and is therefore an index of *hierarchical structure*. We used the weighted rich-club coefficient [18]: for each degree threshold $k$, it compares the total connection weight among nodes of degree greater than $k$ with the largest weight those edges could carry if the strongest connections of the entire network were concentrated among them,

$$\phi^w(k) = \frac{\sum_{i,j\,:\,d_i,d_j>k} A_{ij}}{\sum_{l=1}^{E_{>k}} w_l^{\text{rank}}}, \qquad (11)$$

where $d_i$ is the (binary) degree of node $i$, $E_{>k}$ is the number of edges among the nodes of degree greater than $k$, and $w_1^{\text{rank}} \geq w_2^{\text{rank}} \geq \cdots$ are the edge weights of the whole network ranked in descending order. To distinguish genuine rich-club organization from the level expected by the network's degree sequence alone, $\phi^w(k)$ is normalized by the same coefficient computed on an ensemble of $100$ degree-preserving random reference graphs generated by edge rewiring, $\langle \phi_{\text{rand}}^w(k) \rangle$. We summarize the resulting normalized curve by its mean over the well-sampled range of degree thresholds $\mathcal{K}$, those for which at least 10 nodes remain above threshold, which trims the unstable high-$k$ tail:

$$\phi_{\text{norm}} = \underset{k\in\mathcal{K}}{\text{mean}} \frac{\phi^{w}(k)}{\langle \phi^{w}_{\text{rand}}(k) \rangle}, \qquad (12)$$

with values greater than one indicating a hub backbone that is denser than chance.

To quantify how brain-like an RNN's topology is, we compared its fingerprint $\mathbf{x}^{(r)} = (Q, E, \phi_{\text{norm}})^{\top}$ to the average human fingerprint $\boldsymbol{\mu}$, defined as the mean fingerprint across the individual HCP-YA participants. A plain Euclidean distance is inappropriate here because the three measures vary by very different amounts across healthy brains and are mutually correlated (the strongest pairwise correlation is between modularity and rich club, with $r \approx -0.3$). We therefore used the Mahalanobis distance [19],

$$D_{\text{M}}^{(r)} = \sqrt{(\mathbf{x}^{(r)} - \boldsymbol{\mu})^{\top}\, \boldsymbol{\Sigma}^{-1}\, (\mathbf{x}^{(r)} - \boldsymbol{\mu})}, \qquad (13)$$

where $\boldsymbol{\Sigma}$ is the covariance of the fingerprint across individual HCP-YA participants. Distance is thus expressed in units of human inter-individual variability. To obtain a similarity score from the distance score, we linearly rescaled $D_{\text{M}}$ across the analyzed networks to a similarity $s \in [0,1]$ (arbitrary units; $s = 1$ for the most human-like network, $s = 0$ for the least). This score is the variable used in the regression and enrichment analyses below.

### Supplementary Text 5: Synaptic density data and preprocessing

**RNNs.** For each network the recurrent weight matrix was saved every $40{,}000$ training steps, giving $16$ snapshots spanning the full $6 \times 10^5$-step run. At each snapshot we summarized the network's wiring by the sum of the absolute values of all recurrent weights, i.e., the total recurrent connection strength at that point in training. Stacking the snapshots gives, for every network, a trajectory of total connection strength over training.

**Human data.** No single dataset providing comparable measurements across multiple cortical regions was available, so we compiled one by harmonizing data from [20] for visual cortex and [21] for auditory and prefrontal cortex. These studies provide a mean synaptic density (synapses per $100\,\mu\mathrm{m}^3$) together with its standard error for each age and cortical region, yielding $40$ summary measurements spanning the visual, auditory and prefrontal cortices from the fetal period (0.52 years post-conception) to late adulthood (72 years of age).

To obtain a smooth reference trajectory we min–max normalized the mean densities to $[0,1]$ and fitted a generalized additive model (a smooth function of $\log_{10}$ age). Because each point is a reported group mean rather than a raw observation, we weighted it by the inverse of its squared standard error, the standard approach in meta-analysis, so that precisely estimated means dominate the fit and uncertain ones contribute little. The fitted smooth was highly significant ($edf = 3.45$, $p < 0.0001$, $85.7\%$ of deviance explained).

**Preprocessing.** The human and RNN trajectories were expressed on a common, unit-free amplitude scale: density min–max normalized to $[0,1]$ on each side. This way, their shapes can be compared directly. Their time axes, however, are not yet comparable: human time is measured in years, whereas an RNN has no intrinsic age and its trajectory runs over training steps. A further complication is that the duration of an RNN's training need not correspond to a *full* human lifespan. A given network's trajectory might align only with an early window of development – for instance, the synaptogenesis rise alone – rather than the complete rise-and-prune arc. Which portion of the lifespan each RNN captures is unknown a priori and must itself be inferred.

We therefore tested a range of candidate lifespans. For each candidate from 3 to 75 years (in 3-year steps), the RNN's training-step axis was mapped onto ages from birth to that candidate. The resulting trajectory was smoothed with a generalized additive model on $\log_{10}$ age, and compared with the human reference curve over the same age window. The match was scored on three properties of the two curves: their overall shape (the correlation between them), the timing of the peak relative to the lifespan, and the fraction of density pruned after the peak. The candidate lifespan minimizing the combined distance across the three properties was taken as that network's fitted age: a small fitted age means the trajectory corresponds only to an early developmental window, a large one that it spans a fuller lifespan. Every network thereby received a single fitted age, with substantial variability across the population.

### Supplementary Text 6: Implementation of the compositional generalization test

Compositional generalization can only be tested if particular combinations of primitives are systematically withheld during training; For this reason, while *delay* primitive was trained in all three families of tasks (Reaching, Decision-making, and Match-to-sample), the *anti* primitive was only trained in the Reach and Decision-making families but never in Match-to-sample, and the *context* primitive only in the Reach family, never in Decision-making or Match-to-sample. This resulted in having 12 tasks that were never experienced during training by the RNNs (i.e., the held-out tasks). Because every primitive was itself learned somewhere, solving these tasks does not require acquiring a new operation, only recombining existing ones.

Each held-out task was paired with a familiar reference task that carried the same primitives but came from a family in which that arrangement had been trained. For example, the delayed contextual decision-making task was paired with the delayed contextual go task. The reference therefore matched the held-out task in complexity while being one the network had already learned.

At test, every weight learned during training (input, recurrent and readout) was frozen. The rule input was then extended by a single new channel with its own randomly initialized connections into the hidden layer, and only these new connections were left trainable; a gradient mask prevented any other weight from changing. The network was then optimized to perform a task using only this new input, with Adam (learning rate $10^{-2}$, batch size $32$), the same recurrent noise injected during training, a categorical cross-entropy loss, and gradient-norm clipping at $1$. Performance was evaluated every $100$ optimization steps on $50$ fresh trials; a task was counted as solved at the first evaluation exceeding $90\%$ accuracy, and optimization stopped early once accuracy reached $99\%$. Each task was allotted a budget of $5{,}000$ optimization steps, and tasks not solved within the budget were treated as right-censored at $5{,}000$. The new input was re-initialized independently for every task. Each selected network was probed on all held-out tasks and, as the baseline described above, on matched familiar training tasks.
